# Potent and durable gene modulation in heart and muscle with chemically defined siRNAs

**DOI:** 10.1101/2024.10.01.616183

**Authors:** Hassan H. Fakih, Clemens Lochmann, Rosemary Gagnon, Ashley Summers, Jillian Caiazzi, Julianna E. Buchwald, Qi Tang, Bruktawit Maru, Samuel R. Hildebrand, Mohammad Zain UI Abideen, Raymond C. Furgal, Katherine Y. Gross, Yen Yang, David Cooper, Kathryn R. Monopoli, Dimas Echeverria, JaeHyuck Shim, Ken Yamada, Julia F. Alterman, Anastasia Khvorova

## Abstract

Small interfering RNA (siRNAs) hold immense promise for treating cardiac and muscular diseases, but robust and scalable delivery to these tissues remains a challenge. Recent advances in delivery strategies to muscle include conjugation of biologics (antibody/antibody fragments, peptides), which are currently in clinical development. However, the manufacturing of biologic-siRNA conjugates is a challenging and complex process. By contrast, lipophilic siRNAs are readily chemically synthesized at scale and support sufficient cardiac and skeletal muscle delivery. In this work, we refine siRNA design elements to enhance potency and durability and support clinically relevant silencing in muscle. Applying this strategy for siRNAs targeting myostatin (*MSTN*), a key target in muscle-wasting conditions, we show that a single subcutaneous dose in mice achieved robust and durable silencing (∼80% inhibition up to 6 weeks, ∼30% at 14 weeks). Biweekly dosing resulted in >95% reduction of circulating *MSTN* for half a year, with no observed systemic or target-related toxicity. *MSTN* inhibition resulted in muscle growth and increased lean muscle mass, correlating with improved grip strength. Interestingly, the functional impact on muscle growth and strength significantly outlasts the target silencing, suggesting extended pharmacological effects. Systemic administration was equally efficacious in all muscle groups tested, including skeletal muscle, heart, tongue and diaphragm. The informational nature of the muscle-active chemically defined siRNA scaffold was confirmed by demonstrating muscle and heart efficacy with three additional targets. Our findings pave the way for potent and long-lasting gene modulation in muscle using chemically defined, lipophilic siRNAs, offering a new avenue for treating muscular diseases.

## INTRODUCTION

Oligonucleotide therapeutics (e.g., siRNAs, ASOs, and SSOs) are now regarded as the third class of drugs after small molecules and biologics, as they revolutionize medicine by enabling potent and efficient modulation of gene expression to previously “undruggable” targets with up to 6-12 month duration of effect (1-4). Currently, there are more than 18 approved oligonucleotide therapies, of which six are siRNA-based (1). The clinical success of oligonucleotide therapeutics hinges on their stability and efficient delivery to the tissue of interest (1,5). Five of the approved siRNA drugs are chemically defined and target the liver, owing to robust hepatic delivery by trivalent N-acetylgalactosamine (GalNAc) conjugate (2). Current efforts are focused on enabling delivery to other organs of interest such as the central nervous system, heart, and muscle (1,2).

The ability to robustly deliver and modulate gene expression in muscle tissues is crucial for developing successful and effective therapeutic intervention for cardiac and skeletal muscle diseases (2,5). Local injection of oligonucleotide therapeutics directly into muscle can silence specific genes, but this approach is limited to the area treated, hindering its overall therapeutic impact (6-9). Therefore, recent efforts have focused on achieving systemic delivery of therapeutics to muscles throughout the body, which is more clinically relevant for treating widespread muscle diseases (9). Delivery via biologic conjugates—e.g., antibody/antibody fragment oligonucleotide conjugation (AOC) and peptide-conjugated oligonucleotides—is at the forefront of clinical development (1,5,10-13). Engineered Centyrin proteins that bind specific antigens are showing promise in early clinical trials (14). Strategies for targeted delivery of biologic-siRNA conjugates hold promise, but challenges include complex administration (e.g., hours-long intravenous infusion), developing efficient conjugation methods, ensuring an antibody retains its target recognition ability (epitope sensitivity), preserving siRNA activity, and overcoming large-scale manufacturing difficulties (15,16). These complexities limit the broad applicability and scalability of these approaches.

Meanwhile, lipophilic siRNAs are readily chemically synthesized at scale and have been shown to support extrahepatic delivery, including heart and muscle (5,17,18). Cholesterol, palmitic, and tocopherol-conjugated siRNAs deliver their payload to muscle following systemic injection and efficiently silence multiple targets (e.g., *ALK4, DMPK, MSTN*) (18,19). However, meaningful silencing required high-dose (>50 mg/kg) with repeated administration and resulted in observed toxicity, limiting tolerability and clinical translatability (19).

Recently, our lab has shown that conjugating the fatty acid docosanoic acid (DCA) to siRNAs (DCA-siRNA) supports efficient heart and muscle delivery (20) with better safety profile, but the observed efficacy was not sufficient to enable clinical translation.

In this work, we explore if systematic engineering of the muscle-delivering scaffold (DCA), including hyper-functional siRNA sequence, chemical modification patterns and novel backbone stabilization can help to achieve robust silencing and durability. We identify a muscle-active scaffold that enables durable and clinically relevant silencing in muscle. Targeting myostatin (*Mstn*), a key gene that promotes muscle-wasting, we show that a single subcutaneous injection in mice achieves robust and durable silencing (∼80% inhibition up to 6 weeks, ∼30% at 14 weeks). Repetitive dosing supported >95% reduction of circulating MSTN protein for half a year with no observable toxicity. MSTN suppression led to increased muscle growth and lean muscle mass that correlated with increased function (grip strength). Interestingly, a single dose of muscle-optimized DCA-siRNA targeting *Mstn* produced comparable muscle growth and strength improvements to that of sustained silencing. Our findings suggests that a single treatment of muscle-optimized DCA-siRNA targeting *Mstn* could provide substantial clinical benefit.. Our findings pave the way for effective and long-lasting gene modulation in muscle using chemically defined, lipid-conjugated siRNAs, presenting a promising new approach for treating muscular diseases.

## METHODS

### Designing siRNAs against myostatin

siRNAs were designed based on an algorithm developed by Shmushkovich et al., (21). Briefly, siRNAs were selected based on a 20-nucleotide target sequence from human (accession number: NM_005259) and mouse (NM_010834.3) *MSTN* transcripts. siRNA sequences were excluded if they had: (a) > 56% of GC content; (b) single-nucleotide stretches of four or more; or (c) perfect homology to human miRNA seeds at position 2–7 of the antisense strand56. To minimize off-target effects, siRNAs were excluded if position 2–17 of the antisense strand had full complementarity to non-target mRNAs. Targeting sequences (including +10/−15 flanking nucleotides) were scored using a weight matrix and top-scoring sequences in each species were identified. Cross-species targeting was determined based on perfect homology of the 16-nucleotide region within the target sequence (positions 2–17) to the target sequence of the other species. siRNAs were named based on the position in the target transcript from which they were extracted (e.g., human siRNA 2155 was extracted from positions 2155–2174 of NM_005259; mouse siRNA 1928 was extracted from positions 1928-1947). *In vitro* screening oligonucleotides sequences and modifications are provided in **Supplementary Table S1**.

### Synthesis and purification of oligonucleotides

Compounds for *in vitro* screening were synthesized using standard solid-phase phosphoramidite chemistry on a Dr. Oligo 48 high-throughput RNA synthesizer (Biolytic) (22). Two sets of 96 fully chemically modified antisense and sense single strands were synthesized, one set for human and the other for mouse, at 1 µmol-scale. Compounds for *in vivo* injections were made using a MerMade 12 (BioAutomation) synthesizer. Standard RNA 2′-O-methyl, 2′-fluoro modifications were applied to improve siRNA stability (Chemgenes and Hongene). Extended nucleic acid modified RNA (exNA) was used at the termini of *in vivo* compounds (in-house and Hongene). Sense strands of *in vitro* compounds were synthesized on a cholesterol-conjugated solid support (Chemgenes), and the sense strands of *in vivo* compounds were synthesized at 10-40 μmol scale on a docosanoic acid-functionalized controlled pore glass (CPG) (Hongene) (17,20,22). Antisense strands were synthesized on CPG functionalized with a Unylinker (Chemgenes), bis-cyanoethyl-N, N-diisopropyl CED phosphoramidite (Chemgenes) was used to introduce a 5′-mono-phosphate for *in vitro* compounds, and custom 5′-(E)-vinylphosphonate phosphoramidite (Hongene) was applied for *in vivo* studies.

All sense strands had a dT_2_ spacer between the oligonucleotide and the conjugate. All strands were cleaved and deprotected using 28% aqueous ammonium hydroxide solution for 20 hours at 55°C, followed by drying under vacuum at 40°C, and resuspension in Millipore H_2_O. Oligonucleotides were purified using an Agilent Prostar System (Agilent Technologies) on a C18 column for lipid-conjugated sense strands and an ion-exchange column for antisense strands. Purified oligonucleotides were desalted by size-exclusion chromatography and characterized by LC-MS analysis on an Agilent 6530 accurate-mass quadrupole time-of-flight (Q-TOF) LC/MS (Agilent Technologies).

The sequences and modifications of *in vivo* oligonucleotides are shown in **Supplementary Table S2**. JAK1 sequences were acquired from the work of Tang *et. al*. and Tang, Gross, Fakih *et. al*., (22,23). MECP2 targeting sequences were acquired from the work by Khvorova and Hariharan (24). HTT targeting sequences were the same as previously reported (25,26).

### Duplex formation

Equimolar amounts of antisense and sense strands were prepared in 1x PBS and incubated at 95°C for 5 minutes, cooled to room temperature and incubated overnight at 4°C. To validate efficient duplex formation, 20 pmol of duplex was loaded onto a non-denaturing 20% tris-borate-EDTA (TBE) gel (Invitrogen # EC63155BOX) and run at constant 180V for 1 hour. The gel was then washed with deionized water for 10 minutes, stained with 1X SYBR Gold Nucleic Acid Gel Stain (Invitrogen # S11494) for 10 minutes and washed again. Bands were visualized on the ChemiDoc MP Imaging system (Bio-Rad # 17001402).

### Cell culture

Human SJCRh30 cells (ATCC # CRL-2061) were cultured in T75 tissue-culture treated flasks (Corning # 353107) containing RPMI-1640 (Gibco) supplemented with 10% fetal bovine serum (Gibco) at 37°C and 5% CO2. Murine C2C12 cells (ATCC # CRL-1772) were propagated in DMEM (Gibco) containing 10% fetal bovine serum under the same conditions. All cells were maintained at 10% – 40% confluency and split every 3 – 5 days until passage number reached 20 or 10, respectively. The supernatant medium was discarded, the cells were washed with DPBS (Gibco) and then detached by incubation with TrypLE Express (Gibco) for 5 min at 37°C and 5% CO2. Trypsinization was stopped by adding fresh medium. Counting of cells was performed by trypan blue staining using a Cellometer Auto 1000 instrument (Nexcelom).

### Primary screening

Screening for *MSTN* and *Mstn* targeting siRNAs was conducted in SJCRh30 cells for human-targeting siRNAs, and in C2C12 cells for mouse-targeting siRNAs. siRNA uptake was viaa passive cellular uptake. Samples were diluted in OptiMEM (Gibco) to 2-fold final concentration. Cells were detached and counted and subsequently, resuspended in media containing 6% fetal bovine serum. 50 µL of cell suspension (7,500 cells for SJCRh30, 7,500 cells for C2C12) were plated with 50 µL of siRNA in a 96-well tissue-culture plate to obtain a final serum concentration of 3% and an siRNA concentration of 1.5 µM (screening) or the corresponding target concentration (concentration-response). For each compound three technical replicates were prepared and incubated at 37°C and 5% CO2. After 72h for SJCRh30 or 144h for C2C12, mRNA was quantified as described below.

### mRNA quantification

mRNA expression was quantified with the QuantiGene Singleplex Assay Kit (Invitrogen # QS0016), a hybridization-based assay, as previously described (20,22). Cultured cells were lysed in a total volume of 250 µL diluted lysis mixture consisting of one part lysis mixture, two parts deionized H2O and 0.167 mg/mL proteinase K (Invitrogen # AM2548). Cell lysates were thoroughly mixed by pipetting up and down 15 -20 times. Probe set validation was performed for each cell line and tissue to determine an appropriate lysate volume that would allow unambiguous and linear mRNA detection. The corresponding volumes were transferred to the capture plate as indicated in the manufacturer’s protocol (Invitrogen # MAN0018628), and the protocol was followed further. Luminescence detection was performed using the SpectroMax M5 Microplate Reader (Avantor).

For mouse tissue lysate preparation and mRNA quantification, tissues were collected at the indicated time points and stored in RNA later solution (Sigma-Aldrich; #R0901) at 4 °C overnight. Punch biopsies from the specified tissues were collected (1×4mm punches) and added to 400 μL of homogenizing solution (Invitrogen; QS0517) containing 0.2 mg/mL proteinase K (Invitrogen, #AM3546), in a QIAGEN collection microtube holding a 3-mm tungsten bead. The tissues were then homogenized for 10 min under 30 Hz of frequency using a QIAGEN TissueLyser II, followed by incubating at 55 °C for 30 min. Lysate was then used as described above for mRNA quantification, or for siRNA accumulation (described below) or stored at -20 °C.

### Animal experiments

All animal procedures were conducted according to the Institutional Animal Care and Use Committee (IACUC) protocols of the University of Massachusetts Chan Medical School (IACUC protocol 202000010) and in accordance with the National Institutes of Health Guide for the Care and Use of Laboratory Animals. FVB female mice (7-9 weeks old) were used for all studies. The colonies were maintained and housed at pathogen-free animal facilities at UMass Chan Medical School with 12 h light/12 h dark cycle at controlled temperature (23 ± 1 °C) and humidity (50% ± 20%) with free access to food and water. When indicated, mice body weight was measured using a standard balance. All mice were ear-tagged prior to any study initiation in order to track each mice over the course of the study.

### *In vivo* efficacy

siRNAs formulated in 1x PBS as described above, were always prepared at 200 µM concentration. The siRNAs were then administered (based on the indicated quantities in each experiment) subcutaneously between the shoulder blades of mice. In the multi-dosed groups, the dosage was maintained constant relative to the original weight (e.g., 20 mg/kg was fixed based on the weight on day 0). Animals were then euthanized at the indicated timepoints, and tissues were harvested and processed for mRNA quantification and siRNA accumulation as described above.

### Quantification of Myostatin, Troponin and CK-MB proteins in plasma

Blood samples were collected from control or siRNA injected animals via cheek bleeding, where 20-50 µl was collected on the indicated times (weekly or biweekly). Blood was collected in EDTA coated tubes, and then spun at 10,000 rpm for 10 mins at 4 °C to separate plasma. Plasma was collected and stored at -80 °C until ready to use for quantification. *MSTN* protein was quantified using ELISA kits (Bio-Techne/R&D systems, # DGDF80) as previously described (20), where 3 µl of plasma was used per sample and following standard provided protocol. For Troponin and CK-MB quantification (NBP3-00456, NBP2-75312), 2 µl of plasma was used per sample and then followed by standard provided protocol for each ELISA kit.

### siRNA accumulation quantification

To quantify siRNA accumulated in tissues following systemic administration, PNA hybridization assay was used on the lysate prepared for mRNA quantification, as previously described with slight modification (20,22,27). All samples that were quantified were weighed to calculate siRNA accumulation per milligram of tissue. Briefly, the accumulation quantified using a custom Alexa488-labeled fluorescent PNA oligonucleotide probe (Alexa488-OO-AAUAAGGAAAGAAGAAUCUUA, PNABio) that is fully complementary to the antisense strand. Prepared lysate was used, and Sodium dodecyl sulphate was precipitated from the lysate by adding 50 μL of 3 M potassium chloride followed by centrifugation for 15 min at 5000×g. siRNAs in the clear supernatant were hybridized to the PNA probe under 95 °C followed by slow cool down. Anion-exchanged chromatography was used to analyze the sample mixtures on an Agilent 1260 Infinity quad-pump HPLC with a 1260 FLD fluorescent detector, and quantification was based on compared to standard curves of spike-in siRNA in tissue lysate as previously described (5).

### MRI imaging and analysis

At indicated times for MRI imaging (e.g., pre-dosing, 6, 10, 19 weeks), mice were weighed and then put under anesthesia (2% isoflurane) followed by supine placement in MRI imaging system (Bruker Biospin MRI 7T BioSpec). Mice remained under anesthesia for the duration of imaging, where the left leg (quad and calf) was captured over 3 minutes of imaging for an acceptable resolution. All images were then analyzed using image-j to quantify volume, where the groups/mice were blinded. Briefly, quad and calf areas were highlighted as ROIs and measured over 7 Z-axis slices; volume in mm^3^ was then calculated by summing the areas and multiplying by Z-slice thickness of 0.4mm.

### Lean muscle mass measurement

Lean muscle mass was measured at week 26 post injection using 1H-MRS (Echo Medical System) imaging system by UMass Chan Medical School Metabolic Disease Center. Mice are awake (no anesthesia) when measurements are done to avoid anesthesia-associated complications.

### Grip strength measurement

Grip strength was measured using Grip Strength Meters (harvardapparatus.com) at week 10. Briefly, grip strength using all limbs for each mouse was measured once if the number does not fluctuate, and twice if there was fluctuation followed by averaging. Data was then normalized and plotted as a percentage of the control NTC-DCA-siRNA treated group.

### Blood diagnostics

Blood sampling for blood diagnostics was performed when mice were terminated via cheek bleed. In total, 200 μL of blood was collected in a lithium heparin-coated BD Microtainer tube (BD, #365965) for blood chemistry test. 100 μL of blood was collected in a K2 EDTA-coated bd Microtainer tube (BD, #365974) for a complete blood count (CBC) test. Blood chemistry and CBC diagnostics were conducted by the Diagnostics Laboratory in the Department of Animal Medicine at UMass Chan Medical School. ALT, BUN, platelet count, and immune cell population percentage is shown; remainder of data is available upon request.

### Graphs and statistical analyses

Data were analyzed using GraphPad Prism 10.1.2 software for Windows (GraphPad Software, Inc., San Diego, CA). For each independent mouse experiment, the levels of human or mouse mRNA silencing was normalized to the mean untreated group (for *in vitro* screening) or NTC group (for *in vivo* studies). *In vivo* data were analyzed using a one-way ANOVA with a post hoc Dunnett multiple comparisons when comparing to control group only, or Tukey multiple comparisons when comparing all groups to each other. Asterisks (*) denote gene expression significance relative to NTC or indicated group with bracket (^*^*P* <0.05, ^**^*P* <0.01, ^***^*P* <0.001, ^****^*P* <0.0001), non-significant was not indicated. Graphs are plotted as mean ± standard deviation (except body weight plot which is plotted as mean ± standard error of mean (SEM)).

## RESULTS

### Identification of hyper-functional lead siRNAs targeting murine and human *myostatin* mRNAs

To develop siRNAs that enable potent gene silencing of *myostatin*, we used a custom siRNA design algorithm (28) to identify a panel of 184 siRNAs that target the 5′-untranslated region (5′-UTR), open reading frame (ORF), or 3′-UTR of the mouse *Mstn* (88 siRNAs) or human *MSTN* (96 siRNAs) mRNAs. The siRNAs were synthesized for *in vitro* screening with full chemical modification, including a 3′-cholesterol moiety on the passenger strand for efficient cellular internalization via passive uptake and rapid testing *in vitro*. Sequences and modifications used for these compounds are summarized in Supplementary Table 1. Primary screening in differentiated murine C2C12 and human SJCRh30 cells identified several siRNA sequences that achieve 80-95% silencing (Figure 1). The IC50 values for lead compounds were derived from seven-point dose-response studies (Figure 1). Based on the IC50 values, the newly identified siRNAs *Mstn*^1928^ (IC50 = 26 nM) and *MSTN*^2155^ (IC50 = 75 nM) were the most potent murine and human siRNA leads, respectively. The previously reported sequence (*Mstn*^1192^ / *MSTN*^1194^) was used as a benchmark (18) (20).

**Figure 1.**
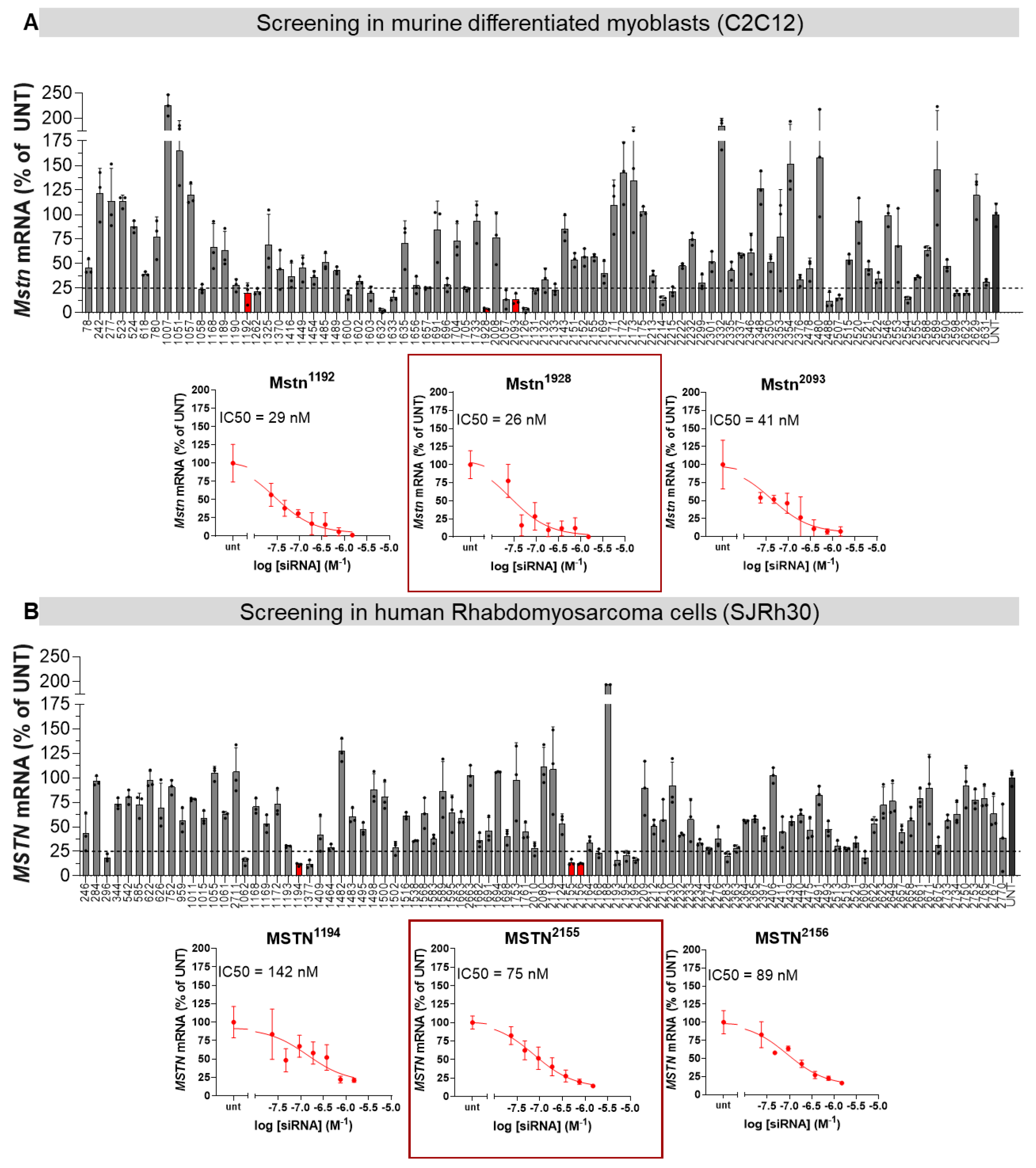
Identification of hyper-functional fully chemically modified siRNAs targeting Myostatin in human and murine cell lines. a panel of 80-96 cholesterol conjugated siRNAs targeting (A) murine *Mstn* gene and (B) human *MSTN* gene were tested for their silencing ability using passive delivery in differentiated murine myoblasts for murine-targeting sequences, and Rhabdomyosarcoma cells for human-targeting sequences. Each screen was followed by measuring the dose-dependent silencing of three selected functional sequences for each species, to obtain the most potent sequence with the lowest IC50. (mRNA measured by the Quantigene 2.0 Assay, n=3, Data represented as mean± S.D.; dotted line on all graphs represents 25% remaining mRNA expression).

### Optimizing guide strand stability, siRNA lead sequence and dosing regimen improves potency in muscle and heart *in vivo*

To compare the *in vivo* activity of the lead *Mstn*^1928^ siRNA sequence to that of the previously reported *Mstn*^1192^ sequence, we synthesized *Mstn*^1928^ in the muscle-delivering DCA-conjugated siRNA scaffold (*Mstn*-DCA-siRNA; Figure 2A). We also included an extended nucleic acid (exNA) modification near the 3′ end of the guide strand (Figure 2A), which significantly increases exonuclease resistance and thereby enhances siRNA accumulation and activity *in vivo*, including in muscle and heart tissues (29). We compared the activities of *Mstn*^1192^-DCA-siRNA with or without exNA and *Mstn*^1928^-DCA-siRNA with exNA (Figure 2B) when injected subcutaneously at 20 mg/kg on days 0 and 3 (20). On day 7, muscle (quadriceps and gastrocnemius) and heart tissues were harvested and processed for *Mstn* mRNA quantification. exNA modification enhanced the silencing activity of *Mstn*^1192^ by ∼50% in all tissues tested compared to without exNA (Figure 2B). exNA modification also improved silencing of a *Jak1* targeting siRNA in quadriceps, consistent with the general ability of exNA to stabilize siRNAs (Supplementary Figure S1).

**Figure 2.**
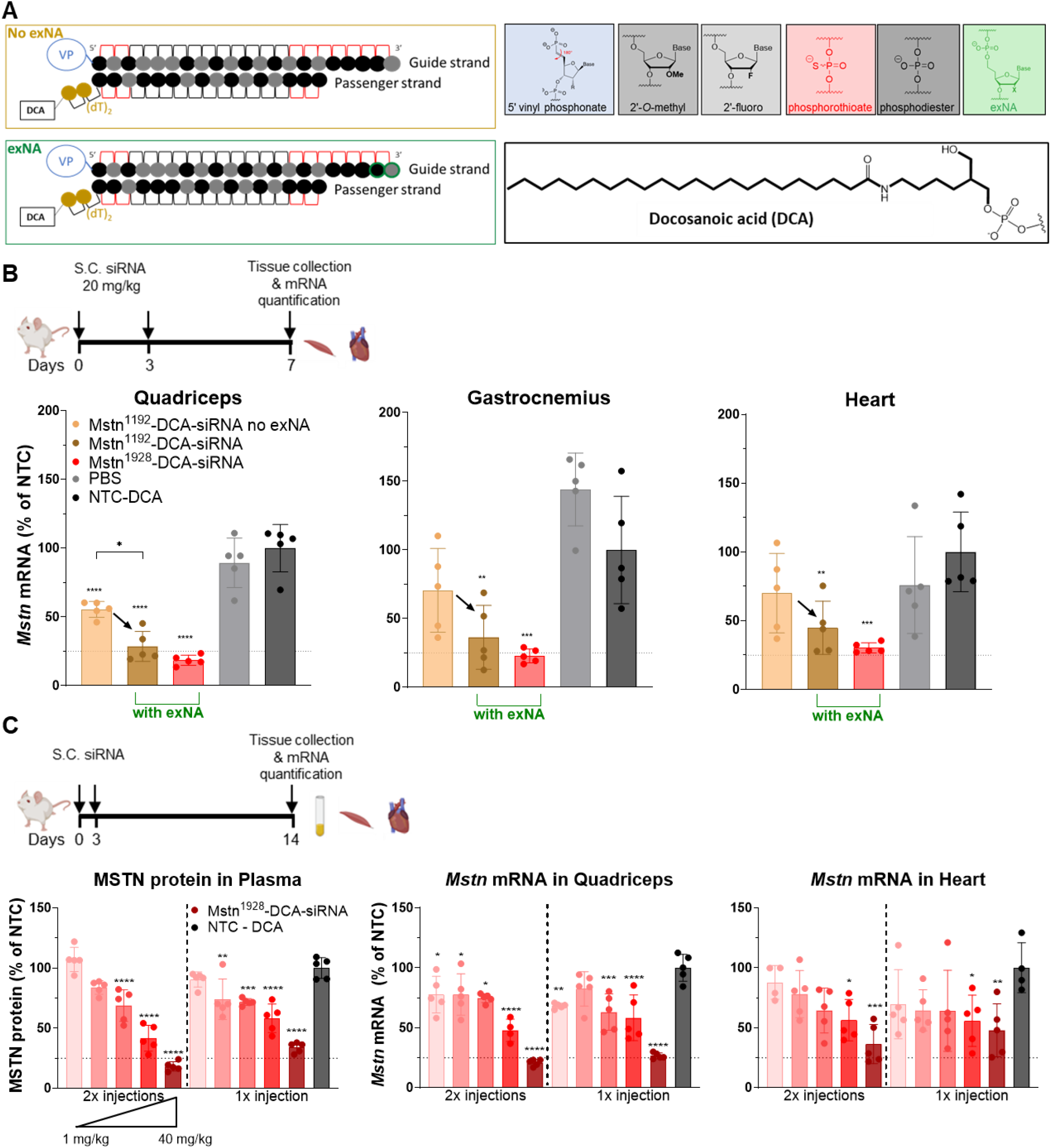
Optimization of siRNA guide strand stability, targeting sequence and dosing regimen improves the potency in muscle and heart. **(A)** Schematic Mstn^1928^-DCA-siRNA chemical modifications used to stabilize the siRNA, with and without exNA terminal modification. **(B)** *Mstn* mRNA silencing in multiple tissues 1-week post injection, using the previously tested Mstn^1192^ sequence with and without exNA inclusion, as well as comparing to the newly identified siRNA lead Mstn^1928^. **(C)** *Mstn* mRNA and MSTN protein silencing 2-weeks post injection at various escalating doses via single and dual injection. All study groups are n=4-5 mice/group. Statistical analysis was performed in GraphPad Prism: for section (B) one-way ANOVA analysis followed by Tukey’s multiple comparisons across all groups, with only significant comparisons against NTC-DCA-siRNA shown, as well as significant comparisons between the treated groups. For section (C) only comparisons to NTC-DCA-siRNA are shown. (* p<0.05, ** p<0.01, *** p<0.001, **** p<0.0001). Subset of data of Mstn1192 from panel B was previously published in Yamada *et. al*. (29). (mRNA measured by the Quantigene 2.0 Assay; protein measured by ELISA; Data represented as mean± S.D.; dotted line on all graphs represents 25% remaining mRNA or protein expression).

Notably, the exNA-modified *Mstn*^1928^-DCA-siRNA showed better potency, achieving 80-90% silencing in skeletal muscle and 75% silencing in the heart, demonstrating the importance of the proper lead siRNA sequence selection.

Focusing on exNA-modified *Mstn*^1928^-DCA-siRNA as a lead, we sought to optimize the dosing regimen and compare a single dose (1, 5, 10, 20, or 40 mg/kg) to two doses (day 0 and 3) impact on efficacy (Figure 2C). At 2 weeks post-injection, we observed a dose-dependent reduction of MSTN protein in plasma and *Mstn* mRNA in quadriceps and heart (Figure 2C). Interestingly, the single 40-mg/kg dose performed slightly better than the 20-mg/kg two-dose regimen (e.g., ∼60% vs 50% reduction in MSTN protein, respectively, p=0.9; and 75% vs 50% reduction of *Mstn* mRNA in quadriceps, respectively, p=0.2), suggesting that a single bolus injection, as long as it is tolerated, is a preferred administration regimen with slight improvement. Repetitive injections produce similar level of mRNA silencing locally in the quadriceps tissue (∼75% reduction *Mstn* mRNA for both 1×40 and 2×40 mg/kg, p=0.2) with some level of improvement on MSTN protein reduction in circulation (∼80% vs 65% reduction in MSTN protein for 2×40 mg/kg vs 1×40 mg/kg, respectively, p=0.9).

### Chemical engineering of modification patterns greatly enhances the durability of silencing effects in muscle and heart tissues

The modification pattern (e.g., 2′OMe-rich vs 2′OMe-2′F balanced) of an siRNA can greatly influence distribution, efficacy, and stability (30,31). We therefore tested different modification patterns in the exNA-modified muscle-targeting scaffold to improve the potency and durable activity in muscle and heart. We synthesized three different patterns (Figure 3A): Pattern 1 has a 2′OMe:2′F ratio of ∼55:45 with a 2′OMe-rich asymmetric tail for increased protection from degradation. This pattern was used in the screen to identify the *Mstn* siRNA lead sequence. Pattern 2 has a 50:50 2′OMe:2′F content, in an alternating pattern, as previously reported (20). Pattern 3 has ∼75% 2′OMe content, which greatly boosts nuclease resistance and stability *in vivo* (2,5,31) (Figure 3A). For each pattern, we generated a version with seven PS (7PS, denoted by hollow circles) modifications and one with four PS (4PS, denoted by filled circles) modifications in the tail of the guide strand. Varying the PS content will determine if more PS groups improve stability and silencing in muscle and heart tissues, even in the presence of exNA modifications (2,31,32).

**Figure 3.**
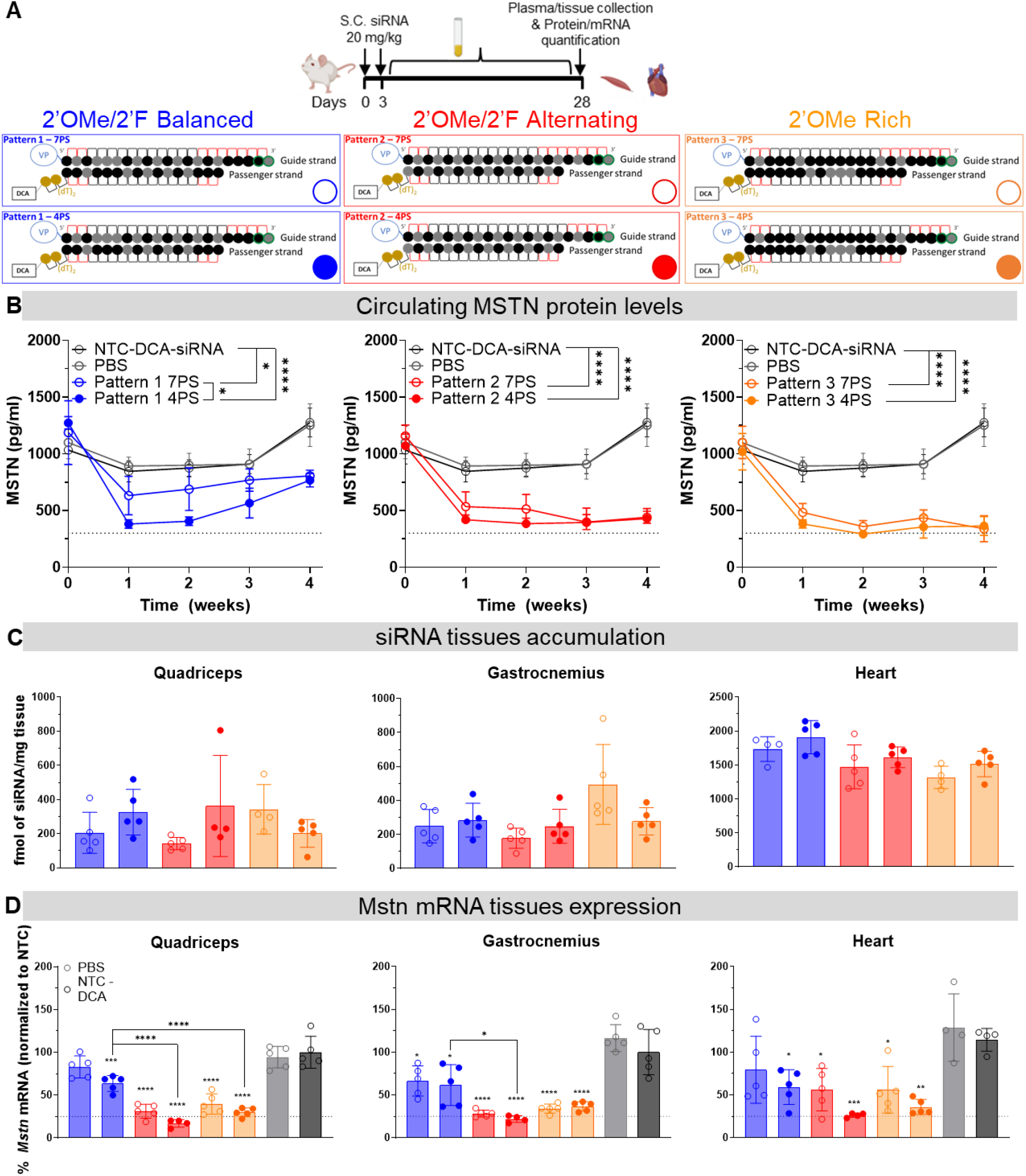
Optimization of the chemical modification pattern and PS tail content optimization greatly enhances durability of silencing in muscle and heart. **(A)** Mstn^1928^-DCA-siRNA with various modification patterns and PS backbone at the 3’ end of guide strand were administered subcutaneously, with plasma being sampled weekly for 4 weeks. **(B)** MSTN protein levels in plasma over time show that the alternating pattern 2 and methyl rich pattern 3 outperform pattern 1 in maximum silencing and durability of consistent silencing over 4 weeks. **(C)** siRNA accumulation in Quadriceps, Gastrocnemius and Heart at 4 weeks post injection show no significant differences in tissue delivery. **(D)** *Mstn* mRNA silencing at 4 weeks is consistent with the MSTN protein levels observed, where pattern 2 with 4PS at the terminus of the guide strand shows the strongest silencing in muscle and heart. All study groups are n=4-5 mice/group. Statistical analysis was performed in GraphPad Prism: for MSTN plasma levels, two-way ANOVA analysis using mixed model was performed, and only significant comparisons are shown; for tissue accumulation and mRNA levels, one-way ANOVA analysis followed by Tukey’s multiple comparisons across all groups, with only significant comparisons against NTC DCA-siRNA shown, as well as significant comparisons between the 4PS groups. (* p<0.05, ** p<0.01, *** p<0.001, **** p<0.0001). (mRNA measured by the Quantigene 2.0 Assay; protein measured by ELISA; Data represented as mean± S.D.; dotted line on all graphs represents 25% remaining mRNA or protein expression).

To compare the activities of each siRNA pattern, we injected mice subcutaneously with two 20-mg/kg doses of each *Mstn*^1928^-DCA-siRNA on days 0 and 3, keeping this dosing regimen for comparative purposes. We sampled plasma weekly for 4 weeks to evaluate kinetics and durability of MSTN silencing (Figure 3A,B). Patterns 2 and 3 both outperformed pattern 1 overall. Pattern 2 and 3, in both PS tail versions, achieved slightly better silencing at week 1 than pattern 1. Whereas patterns 2 and 3 maintained silencing throughout the 4 week analysis, pattern 1 gradually lost silencing activity after week 1.

Interestingly, pattern 1 with 4PS was slightly more active than with 7PS at weeks 2 and 3, before eventually converging on to the same level. We observed no significant difference between 7PS and 4PS in patterns 2 and 3 in terms of durability, except for a slight improvement observed in pattern 2 4PS vs pattern 2 7PS.

At week 4, animals were euthanized, and muscle/heart tissues were collected to measure siRNA accumulation and *Mstn* mRNA knockdown (Figure 3C,D). All three patterns accumulated to similar levels (Figure 3C), suggesting that differences in silencing durability are unrelated to changes in distribution. As for mRNA knockdown, the data correlate with the MSTN protein levels, with patterns 2 and 3 outperforming pattern 1 (regardless of PS version) in each tissue analyzed. Additionally, we observed a statistically significant improvement in silencing activity in patterns 2 and 3 compared to pattern 1, specifically in the 4PS version (statistics for 7PS versions not shown). We also confirmed that MSTN protein levels in quadriceps and gastrocnemius correlate with the mRNA silencing and circulating protein silencing, by achieving 60% protein silencing (Supplementary Figure S2).

Overall, these data show that optimizing the modification pattern—2′OMe:2′F ratio and PS content—improves the potency and durability of our lead Mstn siRNA muscle and heart.

### Potent and durable myostatin silencing produces significant phenotypic changes

For over two decades, myostatin’s role as a major inhibitor of muscle growth has fueled efforts to target it for muscle-wasting diseases like cachexia and aging (33,34). Lowering of myostatin levels induces muscle generation and growth providing for an opportunity to phenotypically characterize the functional consequence of the siRNA-based target modulation in muscle.

To assess the impact of lowering myostatin on phenotypic changes, we treated wild-type mice with exNA-modified *Mstn*^1928^-DCA-siRNA in modification pattern 2 with 4PS. We administered the siRNA subcutaneously once at 10, 20, and 40 mg/kg. We also dosed a group with 40 mg/kg every other week (40mg/kg/2wks) to assess both the impact of completely silencing *MSTN*, as well as the safety profile of repetitive exaggerated lipophilic siRNA (Figure 4A).

**Figure 4.**
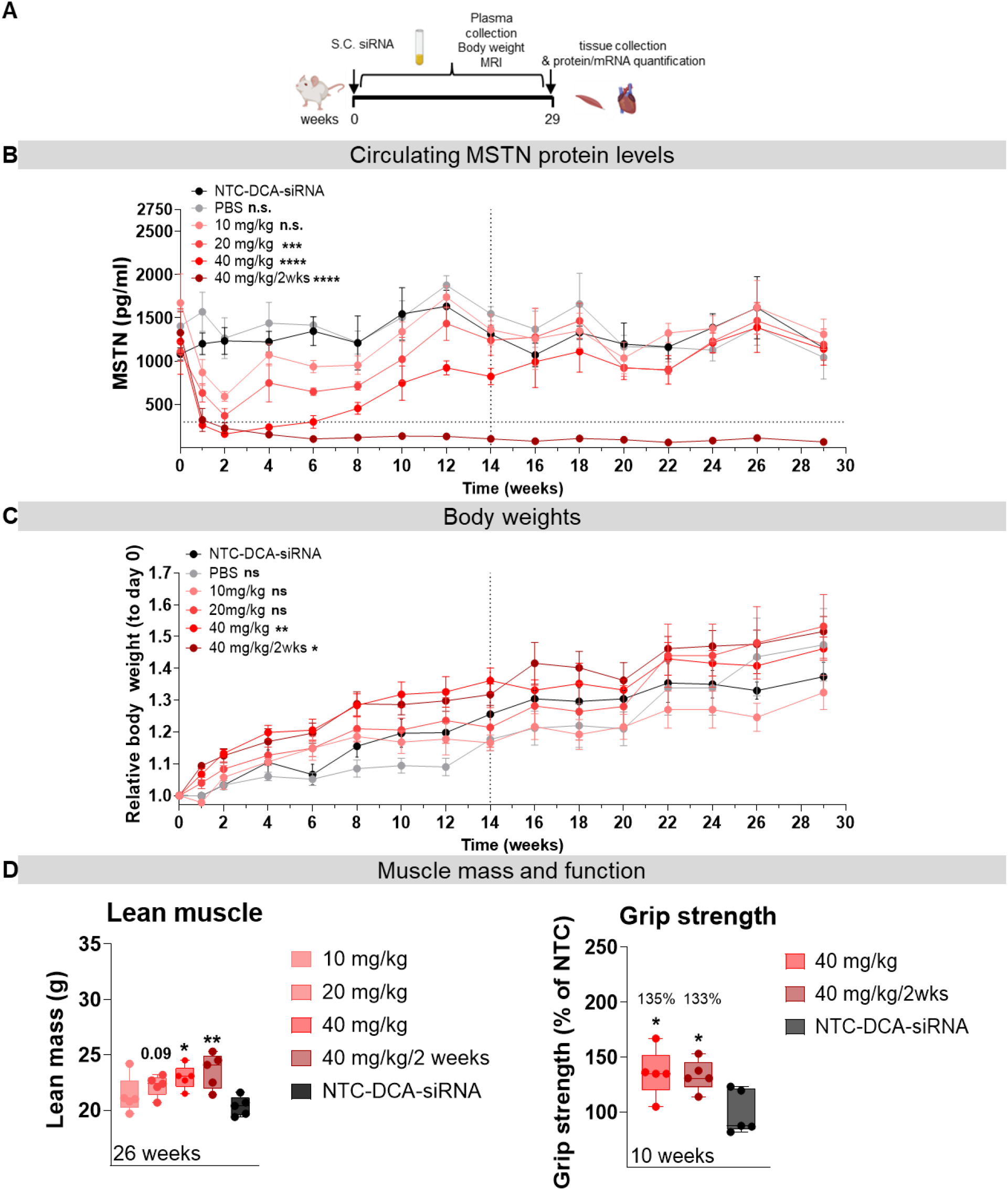
Potent and durable silencing of *Mstn* using the optimized siRNA scaffold leads to prolonged duration of effect and strong phenotypical changes associated with muscle growth. **(A)** Schematic experimental plan, where a Mstn^1928^-DCA-siRNA in Pattern 2 with 4PS and 2 exNA modifications was administered as a single injection at various doses, followed by biweekly plasma collection and body weight measurement, as well as MRI imaging and terminal mRNA quantification for 7 months. **(B)** MSTN plasma levels of different dosing cohorts show significant MSTN protein inhibition as early as one week, with -80% inhibition in the single 40 mg/kg dose up to 6 weeks, lasting up to 14 weeks at ∼30%. Multi-dosed cohort (40mg/kg/2wks) shows near-complete inhibition of MSTN in circulation for half a year. **(C)** Relative body weight increase is observed for all dosing cohorts, with 40 mg/kg and 40 mg/kg/2wks showing the highest increase, significant up to 14 weeks. **(D)** Lean muscle mass composition at 26 weeks post injection shows a dose-dependent increase, with high-dose cohorts showing the biggest gain in lean muscle mass, which translated to increased **(D)** grip strength relative to control group NTC-DCA-siRNA. All study groups are n=4-5 mice/group. Statistical analysis was performed in GraphPad Prism: for MSTN plasma levels and body weights, two-way ANOVA analysis using mixed model was performed, followed by Bonferroni P-value correction, and only significant comparisons are shown. Body weight statistics are analyzed only up to 14 weeks where siRNA is active; lean muscle mass and grip strength one-way ANOVA analysis followed by Tukey’s multiple comparisons across all groups, with only significant comparisons shown. (* p<0.05, ** p<0.01, *** p<0.001, **** p<0.0001).,. (protein measured by ELISA; Data represented as mean± S.D for protein, lean muscle and grip strength and as mean± S.E.M for body weights; dotted y-axis line represents 25% remaining protein expression, dotted x-axis line represents 14 weeks timepoint where body weight statistical analysis is conducted).

Statistically significant silencing was observed at all doses tested with the both potency and durability showing dose dependency. Interestingly, the maximum inihibition was observed at two weeks for all cohorts. Both 20 and 40 mg/kg achieved at leaste 75% silencing at two weeks, with 40 mg/kg dose induced maximum silencing lastin gat least for 6 weeks and partial and significant modulation of myostatin were seen for at least 14 weeks.

The Supplemental Figure S3 shows early silencing kinetics (0.1,3, 7). Compared to the PBS and non-targeting siRNA (NTC-DCA-siRNA) groups, treatment with *Mstn*^1928^-DCA-siRNA reduced MSTN protein by 50% at 3 days post-injection (Supplementary Figure 3) with no detectable sielncign observed at day 1.

The cohort treated with 40-mg/kg/2wks reached a steady-state near complete (>95%) silencing of MSTN at week 6, which was maintained throughout the 29-week long experiment.

*MSTN* inhibition caused a dose-dependent increase in body weight (Figure 4C). The 40 mg/kg and 40 mg/kg/2wks doses increased body weight to similar levels, and the increases were significantly greater than the other cohorts up until 14 weeks (**p<0.01 for 40 mg/kg and *p<0.05 for 40 mg/kg/2wks) after which body weights varied widely in all cohorts (non-significant at 29 weeks for any cohort). As for the 10 and 20 mg/kg doses, near-significant body weight increase was observed up to 6 weeks only.

To determine if the increases in body weight results from muscle gain, we quantified the total lean muscle mass via full body MRI imaging for body composition at 26 weeks post-injection (Figure 4D). We observed a dose-dependent increase in lean muscle mass, with the 40 mg/kg and 40 mg/kg/2wks cohorts showing similar and significant increases (∼13% increase *p<0.05 for 40 mg/kg and ∼15% p*<0.01 for 40 mg/kg/2wks), correlating with the similar increase in body weight.

To assess if the increased muscle mass translates to improved function, we measured the total grip strength of the 40 mg/kg and 40 mg/kg/2wks cohorts at 10 weeks post-injection (Figure 4F). Grip strength was increased by 33-35% in both cohorts compared to that in the NTC-DCA-siRNA control group, indicating that the increased muscle mass is indeed functional and translates to improved strength.

### MRI imaging of quadriceps and gastrocnemius show sustained muscle mass gain

To visually assess changes in muscle mass in mice treated with 40 mg/kg and 40 mg/kg/2wks, we performed a randomized and blinded analysis using MRI to measure changes in the volumes of both quadriceps and calf (gastrocnemius and soleus) at 0, 6, 10 and 19 weeks post-injection (Figure 5A). Regions of interest (ROIs; Figure 5B) were drawn around the quadriceps and calf for each mouse (n=5 per cohort) and a Z-stack of seven images was used to calculate muscle volume at each time point. Notably, we observed significant and similar increases in quadriceps and calf volumes (∼45% and ∼35%, respectively) in the treated cohorts across all time points measured, compared to the pre-injection baseline (week 0) (Figure 5C). For the control NTC-DCA-siRNA group, we only observed a significant change at week 19, due to standard/natural growth. Interestingly, the increase in quadriceps and calf volume was also similar between the 40 mg/kg and 40 mg/kg/2wks groups, in accordance with the body weight, lean muscle and grip strength increases.

**Figure 5.**
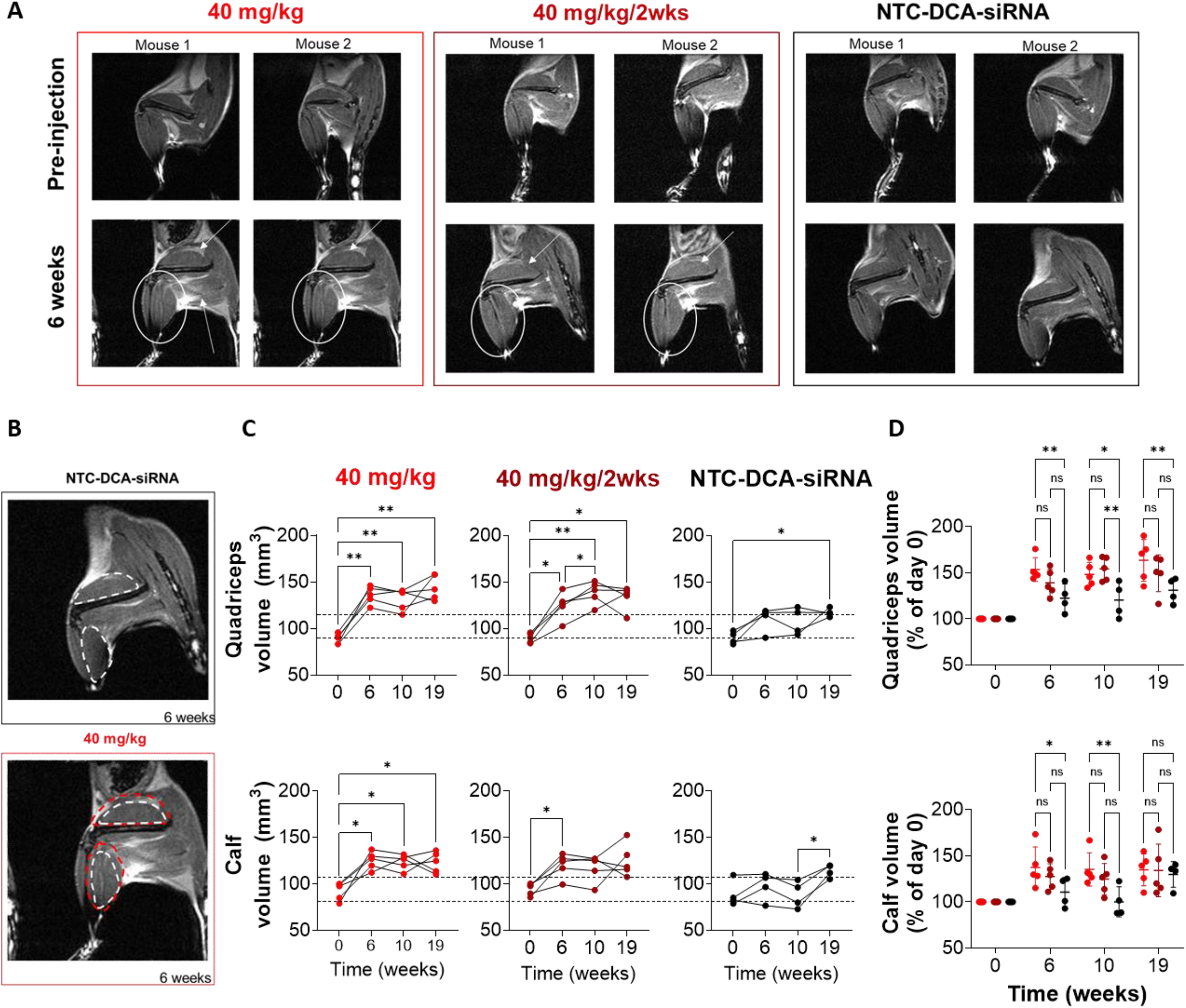
*Mstn* inhibition leads to significant and sustained muscle growth in quadriceps and calf volumes visualized via MRI imaging. **(A)** Representative MRI images of lower left limb of mice from the 40 mg/kg, 40 mg/kg/2wks and NTC-DCA siRNA groups, at pre-injection and 6 weeks timepoint. **(B)** Sample image processing to quantify area of quadriceps and calf, with ROI highlighted. The ROI of quadriceps and calf from NTC in yellow is overlayed on that of the 40 mg/kg sample image to demonstrate increased area and hence volume of corresponding muscles. **(C)** Quadriceps and calf volumes at various time points (0, 6, 10 and 19 weeks) for the imaged groups are quantified blindly and showcased for each individual mouse.**(D)** Normalized quadriceps and calf volumes of each group (to t=0, pre-injection) shows the increase in muscle volume for the treated groups in comparison to control siRNA. All study groups are n=4-5 mice/group. Statistical analysis was performed in GraphPad Prism: row-matched, one-way ANOVA analysis was performed followed by Tukey’s multiple comparisons across all timepoints within each group, with only significant comparisons shown. (* p<0.05, ** p<0.01, *** p<0.001, **** p<0.0001).

By normalizing quadriceps and calf volumes of each group to their initial volume pre-injection (Figure 5C), we observe that mice treated with *Mstn*^1928^-DCA-siRNA at 40 mg/kg have a significant increase in muscle volume at both 6 and 10 weeks compared to NTC-DCA-siRNA treated mice. As for the 40 mg/kg/2wks group, we see a trend in quadriceps volume increase at week 6 and 19, and a statistically significant increase in quadriceps volume at week 10 (p<0.001). Similar trends were observed for the calf volume. Overall, we observe that both single and repetitive treatment with *Mstn*^1928^-DCA-siRNA results in a significant and sustained muscle volume increase relative to NTC-DCA-siRNA.

### Muscle-optimal siRNA scaffold achieves potent silencing in different muscle groups

Although we focused our phenotypic assessments on lower limbs, systemic injection of *Mstn*^1928^-DCA-siRNA is expected to silence *Mstn* in all muscle tissues. Indeed, at 3 weeks after the final injection of 40 mg/kg/2wks *Mstn*^1928^-DCA-siRNA, *Mstn* mRNA was reduced in muscles throughout the body, including lower limb (quadriceps, gastrocnemius), upper limb (biceps), diaphragm, and even tongue.

### Muscle-optimal siRNA scaffold is modular and results in potent silencing when programmed with different sequences and targets

To show that the muscle-optimal scaffold supports silencing of genes other than *Mstn*, we designed muscle-optimal siRNAs (pattern 2 with 4PS and exNA) targeting Janus kinase 1 (*Jak1*; two different sequences), methyl CpG binding protein 2 (*Mecp2*), and huntingtin (*Htt*), for which we have previously identified potent lead sequences (22-24,35). Four weeks post injection of a 40 mg/kg dose, we observed ∼80% silencing of *Jak1* (both sequences), 75% silencing of *Mecp2*, and 50% silencing of *Htt* in the quadriceps and ∼60% silencing of all targets in the heart (Figure 6B). Thus, the muscle-optimized scaffold can be easily reprogrammed to silence any gene of interest in muscle and heart.

**Figure 6.**
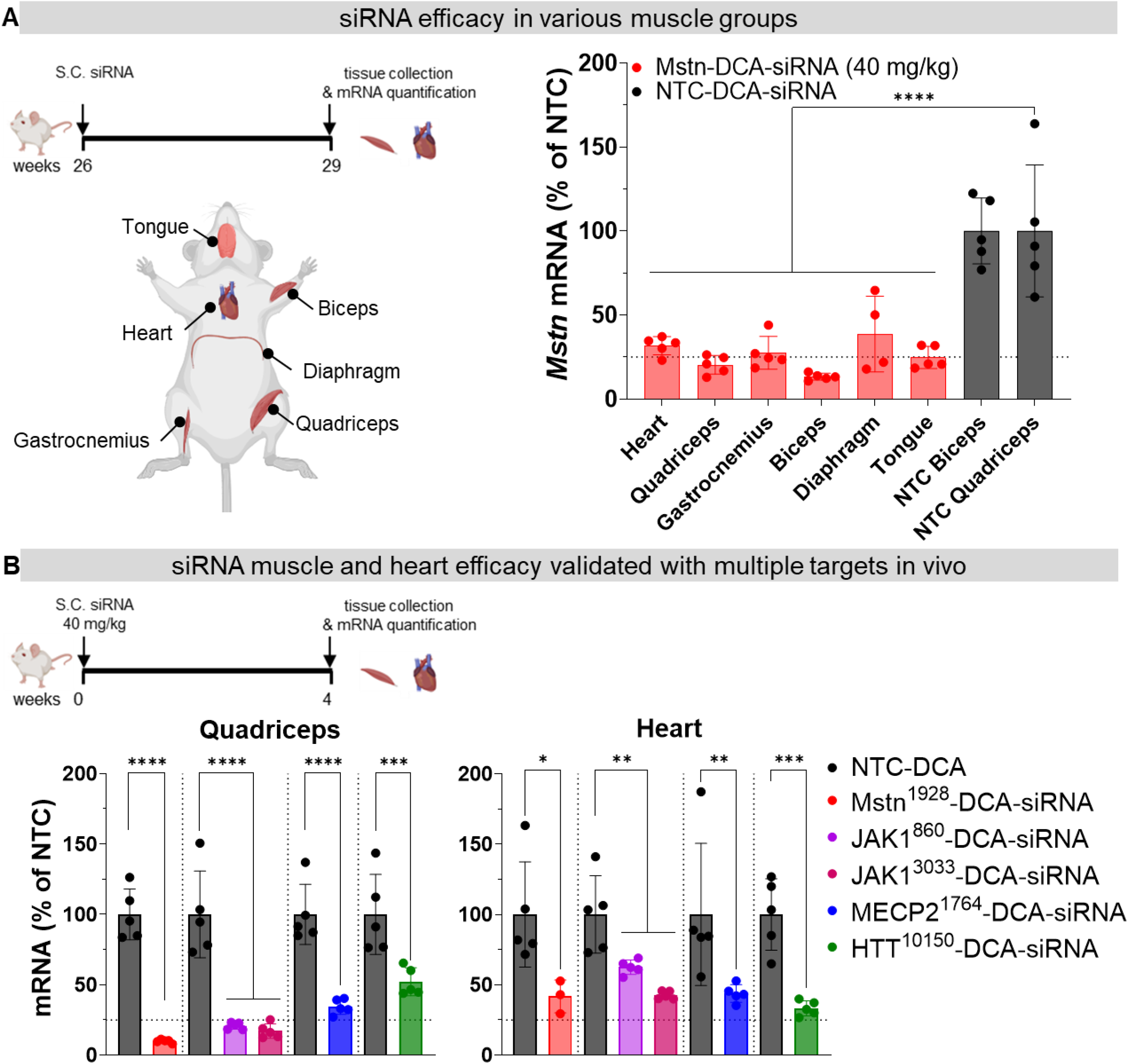
Robust silencing is observed in multiple muscle tissues and for other targets using the optimized DCA-siRNA scaffold. **(A)** Mstn silencing is not limited to lower limb muscle tissues but also observed in upper limb, internal and oral muscles. Tissues were harvested from the 40 mg/kg/2wks cohort 3 weeks post last administration. **(B)** Potent silencing n muscle and heart is not limited to Mstn, as we observe more 75-90% silencing in quadriceps and 50-70% silencing in heart for 3 other targets: Jak1, Mecp2, and Htt. Different targets are separated by dotted vertical lines. All study groups are n=4-5 mice/group. Statistical analysis was performed in GraphPad Prism: one-way ANOVA analysis followed by Tukey’s multiple comparisons across all groups, with only significant comparisons against NTC-DCA-siRNA shown. (* p<0.05, ** p<0.01, *** p<0.001, **** p<0.0001). (mRNA measured by the Quantigene 2.0 Assay; Data represented as mean± S.D.; dotted line on all graphs represents 25% remaining mRNA or protein expression).

### Muscle-optimal siRNA scaffold with exNA shows no detectable toxicity at exaggerated repeat dosing

To assess potential toxicity associated with exaggerated dosing of Mstn^1928^-DCA-siRNA, we performed end-point analyses of 40 mg/kg and 40 mg/kg/2wks cohorts. The 40 mg/kg/2wks cohort received a total cumulative sdose of 10.4 mg (∼ 520 mg/kg) of DCA-siRNA. At week 29, blood was collected for blood chemistry and cell counts analyses, to assess tissue damage and immune reaction (Figure 7A). Compared to the controls, the 40 mg/kg and 40 mg/kg/2wks treatments did not increase liver (ALT) or kidney (BUN) damage markers (Figure 7A), platelet counts, nor immune cells populations. Moreover, we observed no significant increase in troponin and creatine-kinase myocardial band (CK-MB)—known markers of heart damage—at day 3, week 4, or week 12 of the study (Figure 7B; we note the latter two timepoints correspond to 2 weeks after an injection in the 40 mg/kg/2wks cohort). Together, these data indicate that the muscle-optimized scaffold supports safe and effective silencing in muscle and heart.

**Figure 7.**
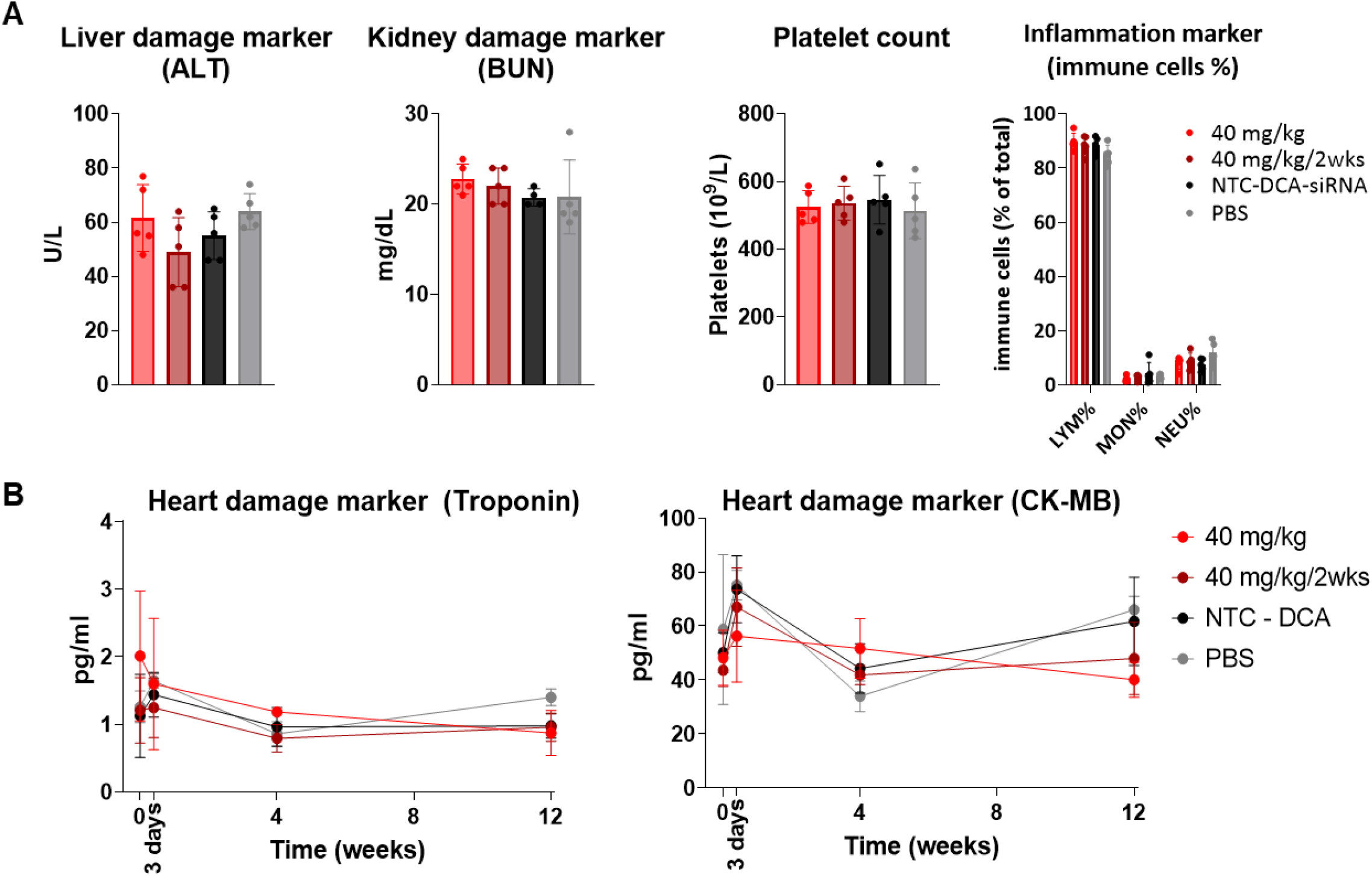
DCA-siRNA scaffold has no observed toxicity even at high doses. **(A)** No alterations in markers of liver (ALT) and kidney (BUN) damage markers, as well as platelet count and immune cell population from complete blood count and blood chemistry assays, in 40 mg/kg and 40mg/kg/2wks (560mg cumulative dose) cohorts in comparison to controls. **(B)** Troponin and CK-MB levels, gold-standard markers of heart damage, are not altered at multiple timepoints measured (0, 3 days, 4-, 8-, and 12-weeks post injection) following *Mstn* silencing, indicating the absence of cardiac damage. All study groups are n=4-5 mice/group.

## DISCUSSION

Multiple technologies are currently being explored to enable the delivery of siRNAs that mediate robust, potent, and durable silencing in muscle tissues (5), including biologic conjugates—e.g., antibodies or peptides—that target specific receptors (1,5,10-13) or lipophilic conjugates that extend circulation time and enable extrahepatic delivery (5,17). Here, we have systematically optimized a simple and readily available lipophilic DCA-conjugated siRNA scaffold and dosing to enhance delivery and activity in muscle and heart tissues. The optimized siRNA targeting the clinically relevant muscle growth inhibitor *MSTN* exhibits superior potency, durability, and desirable phenotypic changes via subcutaneous administration at clinically relevant doses, significantly lower than other modalities. Our findings pave the way for potent and long-lasting gene modulation in muscle using chemically defined, lipid-conjugated siRNAs, offering a new avenue for treating muscular diseases.

### Comparing biologic and lipophilic siRNA conjugation strategies for muscular delivery

Biologic and lipophilic siRNA conjugates offer distinct advantages and disadvantages for muscle delivery (2,16,36). Biologic conjugates, such as antibody-conjugated siRNAs, often require complex and expensive manufacturing processes due to the separate production of the biologic and oligonucleotide components (37). Additionally, the regulatory pathway of biologics under the FDA’s Center for Biologics Evaluation and Research (CEBR FDA) can be more intricate. In contrast, lipophilic conjugates—like the DCA scaffold—are chemically defined and synthesized in a single step, streamlining production (36), and the regulatory pathway under the FDA’s Center for Drug Evaluation and Research (CEDR FDA) regulatory pathway for these chemically defined conjugates is generally less complex.

Stability, dosing, and administration are also crucial factors to consider. Biologic conjugates typically have low drug substance content (10-30%) (38) and are labile at elevated temperature (37). Lipophilic conjugates, however, offer high drug substance content (95%) and superior stability, which simplifies storage and dissemination. But the hydrophobicity of lipophilic conjugates may lead to aggregation and dosing challenges and long-term safety profile of repetitive administration is not established. In terms of dosing, the low solubility and high mass of biologic-conjugated siRNAs requires hundreds of milliliter formulation and hours-long intravenous infusions in clinic (38,39), whereas lipophilic siRNAs are predicted to require a much smaller clinical volume of 5 ml, which can be administered both subcutaneously (divided dose) or intravenously (within minutes). Antibody-oligonucleotide conjugates (AOCs) have shown enhanced delivery and potency in preclinical and clinical studies (12,14,38,40), including better muscle-to-liver distribution compared to DCA-siRNA. For instance, a Transferrin-AOC achieved a 1:5 muscle-to-liver ratio and 1:2 heart-to-liver ratio (12), whereas DCA-siRNA distribution was ∼1:10 muscle-to-liver. Transferrin-AOC also achieved maximum silencing at lower doses than DCA-siRNA (12). Ultimately, the choice between biologic and lipophilic conjugates for muscle delivery will depend on specific clinical needs and considerations.

### Identification of hyper-functional siRNA sequences is crucial for potent durable silencing

Previous siRNA studies achieved ∼50% silencing of *Mstn* in muscle tissues but required high doses (18,20). The siRNA sequence is crucial for overall potency, as it impacts RISC loading, accessibility of the targeted mRNA region, and ultimately, target inhibition(41). Our extensive screen of *Mstn* siRNAs identified lead sequences that enable >90% silencing in cultured cells and >75% silencing in muscle tissue at significantly lower doses than previously reported lipophilic siRNAs (18,20). We believe the enhanced efficacy of the sequence contributes to improved potency and durability. The lead scaffold design exhibits extended silencing (∼80% up to 6-8 weeks) compared to the previous work with a suboptimal sequence (∼45% at 1 month) (18,20).

### Rational chemical stabilization boosts activity and prolongs durability of siRNA in muscle and heart

Chemical modifications, including the 3′-stabilizing exNA modification, improve stability and accumulation in muscle and heart (2) (20) (29). The most common siRNA degradation product is the n-1 metabolite from the 3′ guide strand (guide strand missing the 3′ terminal base) (42). For that, we utilized the recently developed 3′ end stabilizing extended nucleic acid (exNA) modification, that shows improved distribution and efficacy in muscle and heart *in vivo*. Indeed, adding exNA to our siRNA design boosted silencing in muscle tissues.

Beyond stability, modification pattern significantly affects the biodistribution and efficacy of siRNAs in tissues (21,30,31,41,43,44). Although a balanced 2′OMe/2′F modification pattern (e.g., pattern 1) performed well in previous work for various tissues (30,31), we found that a scaffold with alternating 2′OMe/2′F modifications and 4 PS backbone modifications (i.e., pattern 2 4PS) leads to more durable silencing activity in muscle. Further work is needed to determine how pattern 2 enhances silencing in muscle—i.e., if enhanced silencing depends on the tissue or the target.

The durability of silencing is affected by several factors. Some modification patterns might increase durability of silencing by enhancing stability, while others might do so by improving RISC loading or endosomal escape (2). Detailed functional studies are therefore needed to better define the relationship between the chemical structure and performance. This highlights the importance of evaluating not only the absolute activity but also the durability, specifically tailored to the target and tissue of interest.

It is also important to highlight that while having a 7PS backbone modification at the 3′ end of the guide strand provides more stability and expected improved durability, we see the opposite for pattern 1, where 4PS displayed better durability at four weeks. Similar studies conducted for liver-targeting siRNAs have also demonstrated the importance of refining the chemical modification content and pattern to boost activity and durability(2). These findings in muscle tissue underscore the importance of continued investigation into the impact of siRNA modification patterns, especially when targeting new tissues and organs.

### Silencing the therapeutically relevant target myostatin leads to positive phenotypic observations with no adverse events

Silencing myostatin is known to induce muscle growth and improved functionality (e.g., strength), which is a desired phenotype to combat muscle wasting observed in multiple conditions (e.g., muscular dystrophies, ALS, sarcopenia, pancreatic cancer). Clinically evaluated approaches were limited to biologics, which block myostatin’s extracellular signaling by targeting the circulating protein. While demonstrating potent decrease in circulating myostatin, they often faced limitations such as off-target effects and insufficient efficacy due to complex signaling interactions governing muscle growth (34). Developing a siRNA therapy for myostatin inhibition at the intracellular level has not been extensively explored, which is what might be needed to observe clinical efficacy resulting from potent and durable inhibition.

The muscle-optimized siRNA design (pattern 2 4PS with exNA) supported remarkable silencing in mice with a single subcutaneous dose. A single 40 mg/kg dose resulted in robust and long-lasting Mstn inhibition, with >80% suppression for 6 weeks and >30% suppression out to 14 weeks. To our knowledge, this represents the most potent and durable suppression of *Mstn* by siRNA yet reported, especially considering the subcutaneous administration route. Furthermore, the single 40-mg/kg dose is a clinically relevant dose for muscle-focused siRNA development and significantly lower than previously reported (18,19). Maximal silencing of Mstn in muscle was enhanced to >95% and maintained using a repeat-dosing supratherapeutic regimen (40 mg/kg/2wks). Notably, silencing *Mstn* either in the single 40-mg/kg dose or the repeat-dosing regimen (40 mg/kg/2wks) led to similar increases in muscle mass and grip strength. While we only quantified the muscle growth in the lower limbs, our systemic siRNA delivery approach was shown to achieve potent *Mstn* mRNA suppression across various muscle tissues following subcutaneous administration. This is particularly advantageous for muscle-wasting diseases, where muscle loss often occurs throughout the body. Similarly, our potent scaffold confers improved activity to other targets as demonstrated by three other genes (*Jak1, Mecp2, Htt*). These findings suggest the broad potential of this approach for therapeutic interventions in muscle and heart diseases requiring potent gene silencing.

### Phenotypic changes of myostatin silencing extend beyond gene modulation levels

We were surprised that a single, 40 mg/kg dose and repeated 40 mg/kg/2wks dosing had similar effects on muscle growth and grip strength. By 19 weeks, although MSTN protein levels had recovered in the single-dose cohort but remained suppressed (>95%) in the repeat-dosing cohort. Despite this difference, both cohorts showed nearly identical increases in quadricep and calf muscle volumes. Moreover, both groups showed similar increases in total lean muscle mass by 26 weeks—near the study endpoint. These observations suggest that muscle growth induced by a single treatment may offer long-term clinical benefit. This possibility might be an improvement over the weekly or monthly administration of MSTN-neutralizing antibody therapies, which have ultimately failed in clinical trials (34,45,46). Future work is needed to determine a long-term dosing schedule that provides the greatest clinical benefit. This raises the possibility of a single dose every 6-12 months being sufficient for therapeutic effect in a clinical setting.

Furthermore, lean muscle mass gain was observed (although not statistically significant) even at the lower 20 mg/kg dose at 26 weeks, compared to the control group (Figure 4D). This suggests that even modest modulation of *Mstn* mRNA in tissues and circulation can lead to positive phenotypic outcomes.

### DCA siRNA is well tolerated with no toxicity observed from neither on-target effects nor general siRNA chemistry at high doses

Although a biweekly 40 mg/kg dose exceeds anticipated therapeutic need, it allowed us to evaluate two key safety aspects: complete target inhibition and potential toxicity associated with the DCA-siRNA chemistry. Although mice received a large cumulative dose of siRNA (560 mg/kg), we found no evidence of liver or kidney damage, changes in immune cell populations, nor cardiac hypertrophy or injury. The luck of observable gross phenotypes was encouraging and supports safety profile of an siRNA-based therapeutic for muscle diseases, including those necessitating high and repeat dosages. Detailed further toxicology studies would be necessary for a complete assessment.

Overall, our findings pave the way for potent long-lasting and safe gene modulation in muscle and heart using chemically defined, lipid-conjugated siRNAs, offering a new avenue for treating muscular diseases.

## CONTRIBUTIONS

HHF, CL, JFA and AK conceived the project. HHF, KY, JFA and AK contributed to the experimental design. HHF. CL, RG, AS, JC, JEB, QT, BM, MZA, RCF, KYG contributed experimentally. HHF synthesized all *in vivo* siRNA compounds, DC synthesized the screening compounds. HHF wrote the manuscript. All authors provided feedback and approved the manuscript.

## CORRESPONDING AUTHOR

Julia F. Alterman and Anastasia Khvorova.

## DATA AVAILABILITY

All data presented in the main text and the supplementary information are available from the corresponding authors upon request.

## ACKNOWLEDGEMENTS

We thank Khvorova lab members (past and present) for insightful discussions and support. We thank the animal medicine core, as well as the Advanced MRI Center (Dr. Robert King) at UMass Chan Medical School, for their help and support on multiple studies. All figures were created with PowerPoint and BioRender.com. We thank Darryl Conte for his thoughtful comments on manuscript preparation.

## SUPPLEMENTARY INFORMATION

Supplementary Figures S1– 3 and Supplementary Tables S1 and S2.

## REFERENCES

1. (2024) Nucleic Acid Therapeutics: Successes, Milestones, and Upcoming Innovation. Nucleic Acid Therapeutics, 34, 52–72.

2. Egli, M. and Manoharan, M. (2023) Chemistry, structure and function of approved oligonucleotide therapeutics. Nucleic Acids Research, 51, 2529–2573.

3. Belgrad, J., Fakih, H.H. and Khvorova, A. (2024) Nucleic Acid Therapeutics: Successes, Milestones, and Upcoming Innovation. Nucleic Acid Therapeutics, 34, 52–72.

4. Ray, K.K., Wright, R.S., Kallend, D., Koenig, W., Leiter, L.A., Raal, F.J., Bisch, J.A., Richardson, T., Jaros, M. and Wijngaard, P.L. (2020) Two phase 3 trials of inclisiran in patients with elevated LDL cholesterol. New England journal of medicine, 382, 1507–1519.

5. Tang, Q. and Khvorova, A. (2024) RNAi-based drug design: considerations and future directions. Nature Reviews Drug Discovery, 23, 341–364.

6. Gebski, B.L., Mann, C.J., Fletcher, S. and Wilton, S.D. (2003) Morpholino antisense oligonucleotide induced dystrophin exon 23 skipping in mdx mouse muscle. Human molecular genetics, 12, 1801–1811.

7. Hagstrom, J.E., Hegge, J., Zhang, G., Noble, M., Budker, V., Lewis, D.L., Herweijer, H. and Wolff, J.A. (2004) A facile nonviral method for delivering genes and siRNAs to skeletal muscle of mammalian limbs. Molecular therapy, 10, 386–398.

8. Laws, N., Cornford-Nairn, R.A., Irwin, N., Johnsen, R., Fletcher, S., Wilton, S.D. and Hoey, A.J. (2008) Long-term administration of antisense oligonucleotides into the paraspinal muscles of mdx mice reduces kyphosis. Journal of Applied Physiology, 105, 662–668.

9. Nicholson, T.A., Sagmeister, M., Wijesinghe, S.N., Farah, H., Hardy, R.S. and Jones, S.W. (2023) Oligonucleotide Therapeutics for Age-Related Musculoskeletal Disorders: Successes and Challenges. Pharmaceutics, 15, 237.

10. Mullard, A. (2022) Antibody-oligonucleotide conjugates enter the clinic. Nat Rev Drug Discov, 21, 6–8.

11. Barker, S.J., Thayer, M.B., Kim, C., Tatarakis, D., Simon, M., Dial, R.L., Nilewski, L., Wells, R.C., Zhou, Y. and Afetian, M. (2023) Targeting transferrin receptor to transport antisense oligonucleotides across the blood-brain barrier. BioRxiv, 2023.2004. 2025.538145.

12. Malecova, B., Burke, R.S., Cochran, M., Hood, M.D., Johns, R., Kovach, P.R., Doppalapudi, V.R., Erdogan, G., Arias, J.D. and Darimont, B. (2023) Targeted tissue delivery of RNA therapeutics using antibody–oligonucleotide conjugates (AOCs). Nucleic Acids Research, 51, 5901–5910.

13. Desjardins, C.A., Yao, M., Hall, J., O’Donnell, E., Venkatesan, R., Spring, S., Wen, A., Hsia, N., Shen, P. and Russo, R. (2022) Enhanced exon skipping and prolonged dystrophin restoration achieved by TfR1-targeted delivery of antisense oligonucleotide using FORCE conjugation in mdx mice. Nucleic Acids Research, 50, 11401–11414.

14. Klein, D., Goldberg, S., Theile, C.S., Dambra, R., Haskell, K., Kuhar, E., Lin, T., Parmar, R., Manoharan, M. and Richter, M. (2021) Centyrin ligands for extrahepatic delivery of siRNA. Molecular Therapy, 29, 2053–2066.

15. Nanna, A.R., Kel’in, A.V., Theile, C., Pierson, J.M., Voo, Z.X., Garg, A., Nair, J.K., Maier, M.A., Fitzgerald, K. and Rader, C. (2020) Generation and validation of structurally defined antibody–siRNA conjugates. Nucleic Acids Research, 48, 5281–5293.

16. Cao, W., Li, R., Pei, X., Chai, M., Sun, L., Huang, Y., Wang, J., Barth, S., Yu, F. and He, H. (2022) Antibody–siRNA conjugates (ARC): Emerging siRNA drug formulation. Medicine in Drug Discovery, 15, 100128.

17. Osborn, M.F., Coles, A.H., Biscans, A., Haraszti, R.A., Roux, L., Davis, S., Ly, S., Echeverria, D., Hassler, M.R. and Godinho, B.M. (2019) Hydrophobicity drives the systemic distribution of lipid-conjugated siRNAs via lipid transport pathways. Nucleic acids research, 47, 1070–1081.

18. Khan, T., Weber, H., DiMuzio, J., Matter, A., Dogdas, B., Shah, T., Thankappan, A., Disa, J., Jadhav, V., Lubbers, L. et al. (2016) Silencing Myostatin Using Cholesterol-conjugated siRNAs Induces Muscle Growth. Molecular Therapy - Nucleic Acids, 5.

19. Østergaard, M.E., Jackson, M., Low, A., E. Chappell, A.G. Lee, R., Peralta, R.Q., Yu, J., Kinberger, G.A., Dan, A., Carty, R. et al. (2019) Conjugation of hydrophobic moieties enhances potency of antisense oligonucleotides in the muscle of rodents and non-human primates. Nucleic Acids Research, 47, 6045–6058.

20. Biscans, A., Caiazzi, J., McHugh, N., Hariharan, V., Muhuri, M. and Khvorova, A. (2021) Docosanoic acid conjugation to siRNA enables functional and safe delivery to skeletal and cardiac muscles. Mol Ther, 29, 1382–1394.

21. Shmushkovich, T., Monopoli, K.R., Homsy, D., Leyfer, D., Betancur-Boissel, M., Khvorova, A. and Wolfson, A.D. (2018) Functional features defining the efficacy of cholesterol-conjugated, self-deliverable, chemically modified siRNAs. Nucleic Acids Research, 46, 10905–10916.

22. Tang, Q., Fakih, H.H., Zain Ui Abideen, M., Hildebrand, S.R., Afshari, K., Gross, K.Y., Sousa, J., Maebius, A.S., Bartholdy, C., Søgaard, P.P. et al. (2023) Rational design of a JAK1-selective siRNA inhibitor for the modulation of autoimmunity in the skin. Nature Communications, 14, 7099.

23. Tang, Q., Gross, K.Y., Fakih, H.H., Jackson, S.O., Zain U.I. Abideen, M., Monopoli, K.R., Blanchard, C., Bouix-Peter, C., Portal, T., Harris, J.E. et al. (2024) Multispecies-targeting siRNAs for the modulation of JAK1 in the skin. Molecular Therapy - Nucleic Acids, 35.

24. Anastasia Khvorova, V.N.H. (2023) Worldwide applications - United States.

25. Fakih, H.H., Tang, Q., Summers, A., Shin, M., Buchwald, J.E., Gagnon, R., Hariharan, V.N., Echeverria, D., Cooper, D.A., Watts, J.K. et al. (2023) Dendritic amphiphilic siRNA: Selective albumin binding, <em>invivo</em> efficacy and low toxicity. Molecular Therapy - Nucleic Acids, 34.

26. Sarli, S.L., Fakih, H.H., Kelly, K., Devi, G., Rembetsy-Brown Julia M., McEachern Holly R., Ferguson Chantal M., Echeverria, D., Lee, J., Sousa, J. et al. (2024) Quantifying the activity profile of ASO and siRNA conjugates in glioblastoma xenograft tumors in vivo. Nucleic Acids Research, 52, 4799–4817.

27. Godinho, B., Gilbert, J.W., Haraszti, R.A., Coles, A.H., Biscans, A., Roux, L., Nikan, M., Echeverria, D., Hassler, M. and Khvorova, A. (2017) Pharmacokinetic Profiling of Conjugated Therapeutic Oligonucleotides: A High-Throughput Method Based Upon Serial Blood Microsampling Coupled to Peptide Nucleic Acid Hybridization Assay. Nucleic Acid Ther, 27, 323–334.

28. Monopoli, K.R., Korkin, D. and Khvorova, A. (2023) Asymmetric trichotomous partitioning overcomes dataset limitations in building machine learning models for predicting siRNA efficacy. Molecular Therapy - Nucleic Acids, 33, 93–109.

29. Yamada, K., Hariharan, V.N., Caiazzi, J., Miller, R., Ferguson, C.M., Sapp, E., Fakih, H.H., Tang, Q., Yamada, N., Furgal, R.C. et al. (2024) Enhancing siRNA efficacy in vivo with extended nucleic acid backbones. Nature Biotechnology.

30. Ly, S., Echeverria, D., Sousa, J. and Khvorova, A. (2020) Single-Stranded Phosphorothioated Regions Enhance Cellular Uptake of Cholesterol-Conjugated siRNA but Not Silencing Efficacy. Molecular Therapy - Nucleic Acids, 21, 991–1005.

31. Biscans, A., Caiazzi, J., Davis, S., McHugh, N., Sousa, J. and Khvorova, A. (2020) The chemical structure and phosphorothioate content of hydrophobically modified siRNAs impact extrahepatic distribution and efficacy. Nucleic Acids Res, 48, 7665–7680.

32. Yamada, K., Hariharan, V.N., Caiazzi, J., Miller, R., Furguson, C., Sapp, E., Fakih, H., Tan, Q., Yamada, N., Furgal, R.C. et al. (2023) Extended Nucleic Acid (exNA): A Novel, Biologically Compatible Backbone that Significantly Enhances Oligonucleotide Efficacy in vivo. bioRxiv.

33. Suh, J. and Lee, Y.S. (2020) Myostatin Inhibitors: Panacea or Predicament for Musculoskeletal Disorders? J Bone Metab, 27, 151–165.

34. Lee, S.-J., Bhasin, S., Klickstein, L., Krishnan, V. and Rooks, D. (2023) Challenges and Future Prospects of Targeting Myostatin/Activin A Signaling to Treat Diseases of Muscle Loss and Metabolic Dysfunction. The Journals of Gerontology: Series A, 78, 32–37.

35. Alterman, J.F., Godinho, B.M.D.C., Hassler, M.R., Ferguson, C.M., Echeverria, D., Sapp, E., Haraszti, R.A., Coles, A.H., Conroy, F., Miller, R. et al. (2019) A divalent siRNA chemical scaffold for potent and sustained modulation of gene expression throughout the central nervous system. Nature Biotechnology, 37, 884–894.

36. Biscans, A., Coles, A., Haraszti, R., Echeverria, D., Hassler, M., Osborn, M. and Khvorova, A. (2019) Diverse lipid conjugates for functional extra-hepatic siRNA delivery in vivo. Nucleic acids research, 47, 1082–1096.

37. Duerr, C. and Friess, W. (2019) Antibody-drug conjugates-stability and formulation. European Journal of Pharmaceutics and Biopharmaceutics, 139, 168–176.

38. Meglio, M. (2023), Neurology Live, pp. NA.

39. Malecova, B., Burke, R.S., Cochran, M., Hood, M.D., Johns, R., Kovach, P.R., Doppalapudi, V.R., Erdogan, G., Arias, J.D., Darimont, B. et al. (2023) Targeted tissue delivery of RNA therapeutics using antibody–oligonucleotide conjugates (AOCs). Nucleic Acids Research, 51, 5901–5910.

40. Barker, S.J., Thayer, M.B., Kim, C., Tatarakis, D., Simon, M.J., Dial, R., Nilewski, L., Wells, R.C., Zhou, Y., Afetian, M. et al. (2024) Targeting the transferrin receptor to transport antisense oligonucleotides across the mammalian blood-brain barrier. Science Translational Medicine, 16, eadi2245.

41. Foster, D.J., Brown, C.R., Shaikh, S., Trapp, C., Schlegel, M.K., Qian, K., Sehgal, A., Rajeev, K.G., Jadhav, V., Manoharan, M. et al. (2018) Advanced siRNA Designs Further Improve In Vivo Performance of GalNAc-siRNA Conjugates. Mol Ther, 26, 708–717.

42. Li, J., Liu, J., Zhang, X., Clausen, V., Tran, C., Arciprete, M., Wang, Q., Rocca, C., Guan, L.-H., Zhang, G. et al. (2021) Nonclinical Pharmacokinetics and Absorption, Distribution, Metabolism, and Excretion of Givosiran, the First Approved <em>N</em>-Acetylgalactosamine–Conjugated RNA Interference Therapeutic. Drug Metabolism and Disposition, 49, 572–580.

43. Davis, S.M., Hariharan, V.N., Lo, A., Turanov, A.A., Echeverria, D., Sousa, J., McHugh, N., Biscans, A., Alterman, J.F. and Karumanchi, S.A. (2022) Chemical optimization of siRNA for safe and efficient silencing of placental sFLT1. Molecular Therapy-Nucleic Acids, 29, 135–149.

44. Foster, D.J., Brown, C.R., Shaikh, S., Trapp, C., Schlegel, M.K., Qian, K., Sehgal, A., Rajeev, K.G., Jadhav, V., Manoharan, M. et al. (2018) Advanced siRNA Designs Further Improve <em>InVivo</em> Performance of GalNAc-siRNA Conjugates. Molecular Therapy, 26, 708–717.

45. Abati, E., Manini, A., Comi, G.P. and Corti, S. (2022) Inhibition of myostatin and related signaling pathways for the treatment of muscle atrophy in motor neuron diseases. Cell Mol Life Sci, 79, 374.

46. Muntoni, F., Byrne, B.J., McMillan, H.J., Ryan, M.M., Wong, B.L., Dukart, J., Bansal, A., Cosson, V., Dreghici, R., Guridi, M. et al. (2024) The Clinical Development of Taldefgrobep Alfa: An Anti-Myostatin Adnectin for the Treatment of Duchenne Muscular Dystrophy. Neurology and Therapy, 13, 183–219.

